# The first report on the use of a zona pellucida binding assay to compare the effects of European bison (Bison bonasus) epididymal spermatozoa cryopreservation in two different extenders

**DOI:** 10.1101/2023.09.11.557052

**Authors:** Maria Eberhardt, Martina Colombo, Sylwia Prochowska, Gaia C. Luvoni, Wanda Olech, Wojciech Niżański

**Author notes:** Correspondence &.

## Abstract

The wisent (or European bison) Bison bonasus is a species that has gone through a bottleneck, resulting in a narrow gene pool of the current population. Presently, the protection program for this species, in addition to protecting the habitat, is aimed at preserving gene pool. One of its elements is creating a bank of gametes with the intention of using them in constantly developed Assisted Reproductive Techniques in this species. In order to use the potential of the gametes stored in the bank as effectively as possible, it is extremely important to assess their post-freezing quality. Wisent epididymal spermatozoa stored in the bank are evaluated using basic semen analysis methods, the CASA system and flow cytometry. So far, functional tests of spermatozoa have not been introduced to the evaluation protocol of spermatozoa obtained post mortem from European bison. This article, for the first time describes the use of the zona pellucida binding test to assess the quality of cryopreserved wisent epididymal spermatozoa. Due to the close relationship between the species, the Zona binding Assay protocol developed for cattle was used. To exclude the influence of the composition of the extender used for cryopreservation, the spermatozoa were cryopreserved in the Extender based on Tris buffer, egg yolk and glycerol and in the commercial extender Andromed^®^. Regardless of the diluent used, 100% of the oocytes had sperm attached. The average number of spermatozoa attached to the oocyte was 66.26 for the Tris-based diluent group and 27.37 for the Andromed^®^ - frozen group. The advantage of the Tris based extender resulted from ZBA seems to coincide with the results of the basic and advanced semen evaluation. The conducted experiment showed that the ZBA protocol developed for cattle is suitable for the evaluation of wisent epididymal spermatozoa.

## 1. Introduction

Assisted reproductive techniques (ART) are a great achievement not only in the reproduction of livestock or companion animals, but also in the dynamically developing methods of wild species protection programmes [1-2]. However, to be successful in their implementation, it is necessary to thoroughly understand the reproductive physiology of a given species and the characteristics of its gametes. The weakest point, limiting the understanding of reproductive physiology and the development of ART in endangered species, is constricted availability of biological material. As a starting point for research on protected animals, techniques already developed in farm or companion animals are adopted [1]. However, materials, methods, and whole experiments designed for one species require significant revision to become suitable to another, taking into account the availability of the material and the physiology of the animals [1]. The basis for the introduction of assisted reproduction in each species is the collection and cryopreservation of gametes [1-2]. The epididymal spermatozoa are considered as a valuable source of genetic material which usefulness in ART has been proven in numerous species [3-4]. In many cases, obtaining spermatozoa post-mortem from the ep-ididymides due to animal welfare and law regulations is the only way to collect material from wild species, including European bison [3].

The European bison (Bison bonasus Linnaeus, 1758), also known as the wisent, is considered an integral part of native fauna of the European continent [5]. Intensive hunting and poaching, together with environmental degradation resulting from the development of civilization and armed conflicts were the main reasons for the extinction of the natural wisent population. In 1919 in the Białowieska Forest (Poland), the carcass of the last free-living wisent individual was found. The future of the whole species was resting on the shoulders of captive individuals (25 females and the 29 males), and gene pool of the current population carries the genes of merely 12 founders [5]. The decades-long activity aimed at restoring the European bison to its natural habitat has been an undeniable success. However, despite the fact that the number of European bison individuals is already more than 9,000 [6], their narrow initial genetic pool carries a certain risk for its future. For this reason, currently, wisent conservation programs are enriched by preserving of genetic diversity by gametes banking and implementation of ART [2,5,7-8]. There are several reports in the literature on the collection and cryopreservation of wisent epididymal sperm [8-10]. In the cited studies, the effects of spermatozoa cryopreservation were assessed using phase contrast microscope, the CASA system and flow cytometry.

Data from human medicine revealed that basic semen analysis could not be considered as a strictly related predictive factor for pregnancy [11]. Detailed movement parameters evaluated by the CASA system may provide more accurate information about sire fertility potential [4]. Fluorescent staining with flow cytometry is an advanced method that allows to look into the structure of the sperm and thus its functioning [12]. However, the results of the flow cytometry could not decide alone about semen fertility potential.

The stage of spermatozoa - zona pellucida interactions is one of the milestones leading to fertilization which reflects sperm competence [11]. Consequently sperm-zona pellucida binding assay (ZBA) was included in evaluation of the semen prior to conventional in vitro fertilisation (IVF) in human laboratories [11]. The ZBA has also been used in animal reproduction studies. In bovine a positive correlation between the ZBA results and IVF was observed [13]. The use of this test to evaluate the results of sperm cryopreservation has been described, among others, in species such as: dog Canis lupus familiaris [14], cattle Bos taurus [15], buffalo Bubalus bubalis [16] and pig Sus domesticus [17]. To the authors’ best knowledge, there are no reports in the literature about the use of Zona Binding Assay to evaluate wisent epididymal spermatozoa.

The aim of this study was to adapt the ZBA protocol to test the competence of cryo-preserved wisent epididymal spermatozoa. Due to the great value of the oocytes obtained from European bison females, it was decided to develop the protocol with the use of heterologous bovine oocytes. The close relationship of these species and the success in obtaining wisent and cattle hybrids[9] determined this choice. This interspecies approach with the use of bovine oocytes was successfully used in research on other wild animal sperm, such as scimitar - horned oryx Oryx dammah, bottlenose dolphin Tursiops truncatus, or sika deer Cervus nippon and red deer Cervus elaphus [18-20]. To detect possible differences in ZBA resulting from extender composition, spermatozoa samples were split and frozen using a) Tris buffer, hen egg yolk and glycerol-based extender (TEG) or b) Andromed^®^, a commercial medium dedicated to bovine semen. In addition, to validate the ZBA protocol, the obtained results were related to the effects of basic and advanced sperm analysis using the CASA system and flow cytometry.

## 2. Materials and Methods

The spermatozoa were obtained post-mortem from 1 wisent individual from free roaming herd and subjected to planned elimination under conditions regulated by Polish law (The Nature Conservation Act 2004). The bovine oocytes used in zona pellucida binding assay were obtained from ovaries collected in a slaughterhouse. No animal was killed to obtain material for these studies. The Approval of the local ethics committee was not required.

### Reagents

The fluorescent probes: Live/Dead Sperm Viability Kit: SYBR-14, propidium iodide (PI); PNA from Arachis hypogaea Alexa Fluor^®^ 488 conjugate; JC-1; YO-PRO-1; C11-BODIPY581/591; acridine orange (AO) were purchased from Thermo Fisher Scientific Inc., Waltham, MA, USA. Rest of chemicals were bought from Sigma-Aldrich Co., St. Louis, Missouri, USA.

#### Collection of spermatozoa

Immediately after the death of the wisent, the epididymides were separated from the testes. The epididymal tails were cut off and then placed on a Petri dish with 5 ml of TRIS based extender (Tris (2.4⍰g), citric acid (1.4⍰g), glucose (0.8⍰g), penicillin (5000⍰IU) streptomycin (100⍰mg) and distilled water up to 100⍰ml). Subsequently, a series of incisions were made with a scalpel blade aimed to release the sperm into the extender. After collection, samples were placed on the heating table for 5 minutes before further analysis.

#### Initial sperm analysis

Spermatozoa obtained from each epididymal tail were treated as separate samples and subjected to initial semen assessment (concentration, motility, viability and morphology) with the use of phase contrast microscope (Nikon Eclipse E200).

For subjective motility assessment, warm stage was connected to the microscope. Ten microliters of samples were placed on the microscope slide and covered with a cover slip. Both microscope slide and cover slip were warmed before the assessment (×⍰400). Subjective motility was assessed by two independent researchers and the mean value was calculated.

Concentration per unit volume [106 cells/ml] was assessed using the Thoma chamber (×⍰400).

To assess the percentage of live and dead spermatozoa, smears of 10 μL of the samples and 10 μL of the eosin-nigrosin dye were made. Pink-stained spermatozoa were classified as dead. Unstained (white) spermatozoa were classified as live.

For morphology assessment, smears from 10⍰μL of samples were made and left for drying for 24 hours. The next day smears were placed in 96% ethanol solution for 5 minutes fixation. After fixation slides were placed in a 1% water solution of eosin for 3 min. After rinsing in distilled water, the smear was stained for 3 min with gentian pigment (methylene blue 2% w/v, gentian violet 0.75% w/v, glycerol 5% v/v in distilled water). After staining, the smears were rinsed, dried and then evaluated. Two hundred spermatozoa were classified. Proximal droplet, head abnormalities, acrosome abnormalities, midpiece defects, dag-like defect, distal droplet, bent tail, detached head, coiled tail were evaluated. Spermatozoa which were not showing particular defects were classified as normal.

#### Semen freezing

After the initial assessment obtained spermatozoa of each epididymis were divided into two equal parts and cryopreserved with two different methods.

One part of spermatozoa was subjected to cryopreservation in Tris buffer, egg yolk and glycerol based semen extender (TEG) according to the protocol used before in wisent [8-9].

Initially, at room temperature, samples were diluted with Tris based extender (Tris (2.4⍰g), citric acid (1.4⍰g), glucose (0.8⍰g), egg yolk (20% v/v), penicillin (5000⍰IU) streptomycin (100⍰mg) and distilled water up to 100⍰ml) to obtain the final concentration 200⍰×⍰10^6^ /ml. Subsequently, samples were placed in a water bath and put into a refrigerator for cooling down to 5⍰°C. When the samples reached a temperature of 5⍰°C, they were diluted with a chilled extender to the final concentration 160⍰×⍰10^6^ cells/mL. The chilled extender was reached with the glycerol in the amount 6% of the final volume. Samples were left for 90⍰minutes equilibration. After equilibration, samples were loaded into 0.25⍰ml french straws (40⍰×⍰10^6^ spermatozoa per straw). The free straw ends were closed with polyvinyl alcohol (PVA). Filled straws were placed in liquid nitrogen vapours for 15 minutes and then immersed in liquid nitrogen and placed in storage tanks.

The second part of spermatozoa was subjected to cryopreservation in Andromed^®^,a commercially available bovine semen extender, according to a previously described protocol [24]. Spermatozoa were diluted at room temperature with Andromed^®^ up to the final concentration 160⍰×⍰10^6^ sperm/ml. Subsequently, 0.25 ml straws were filled with extended spermatozoa (40⍰×⍰10^6^ spermatozoa per straw). Each free straw end was closed with PVA. Straws were placed in the fridge for slow cooling down to 4°C for 4 hours. After equilibration, straws were placed in liquid nitrogen vapors for 15 minutes. After equilibration time, straws were immersed in the liquid nitrogen and placed in the storage tanks.

#### Thawing

The straws were removed from liquid nitrogen tank and placed for 30 seconds in a water bath (37⍰°C). After thawing, spermatozoa were subjected to evaluation.

#### Post- thawing initial semen assessment

After thawing samples were subjected to the same as described above, initial analyses - subjective motility, morphology and viability.

#### Assessment of thawed sperm movement parameters with CASA

The HTM IVOS version 12.2 (Hamilton-Thorne Biosciences Beverly, MA, USA) semen analyzer was used for detailed movement characterization of thawed spermatozoa. The CASA setups devoted for bull spermatozoa were used (frame rate (60⍰Hz), frames acquired (30), minimum contrast (80), minimum cell size (5 pixels), low VAP cut-off (30⍰μm/sec) and low VSL cut-off (15⍰μm/sec) [25].

The percentage of motility (MOT, %) and progressive motility (PMOT, %), were assessed. Following sperm movement characteristic were evaluated: curvilinear veloc-ity VCL (μm/s), average path velocity VAP (μm/s), straight line velocity VSL (μm/s), linearity LIN (%), straightness STR (%), amplitude of lateral head displacement (ALH, μm), beat cross frequency BCF (Hz). Rapid, medium, slow and static sperm subpopula-tions were also distinguished.

#### Evaluation of thawed spermatozoa by the fluorescent staining and flow cytometry

Membrane and acrosome integrity, lipid peroxidation, apoptosis like changes, mitochondrial activity, membrane lipid disorders and chromatin status were assessed.

All parameters were evaluated using Guava EasyCyte 5 (Merck KGaA, Darmstadt, Germany) cytometer. The fluorescent probes used in the experiment were excited by an argon ion 488⍰nm laser. Gametes acquisitions were analysed with the GuavaSoft™ 3.1.1 software (Merck KGaA, Darmstadt, Germany). The non-sperm events were gated out based on scatter properties and not analysed. A total of 10,000 events were analysed for each sample [8].

The staining used were performed according to protocols routinely used in our laboratory, which have been previously described in wisent [8].

SYBR-14 stain combined with propidium iodide (PI) was used for assessing of sperm membrane integrity. Spermatozoa with intact membranes emit green fluorescence. Cells showing red fluorescence were classified as dead.

Lectin (PNA, Peanut Agglutinin) stain from Arachis hypogaea Alexa Fluor^®^ 488 conjugate combined with propidium iodide (PI) was used for acrosome integrity evaluation. Spermatozoa emitting green fluorescence 427 were classified as those with damaged acrosome.

The cyanine dye JC-1 (5,5⍰,6,6⍰-tetrachloro-1,1⍰,3,3⍰-tetraethylbenzimi-dazolylcar-bocyanine iodide) with PI was used to determine mitochondrial activity. Spermatozoa emitting orange fluorescence were classified as live having high mitochondrial membrane potential and emitting green fluorescence as those live with low mitochondrial potential.

C11-BODIPY581/591(BODIPY, Boron dipyrromethene difluoride) probe combined with PI was used for lipid peroxidation assessment. Spermatozoa which remain un-stained spermatozoa were classified as living population without lipid peroxidation.

YO-PRO-1 (Oxazole yellow, 4-[(3-methyl-1,3-benzoxazol-2(3H)-ylidene)methyl]-1-[3-trimethylammonio)propyl] quinolinium diiodide) dye combined with PI was used to detect spermatozoa with apoptosis like changes. Green fluorescence is characteristic for cells showing apoptotic like changes. Unstained spermatozoa were categorized as living population without apoptotic like changes.

Acridine orange (AO) dye was used to determine the chromatin status of thawed spermatozoa. Spermatozoa with normal DNA configuration were emitting green fluorescence. These gametes emitting red fluorescence classified as cells with denatured DNA.

#### The adaptation of zona pellucida binding assay for wisent frozen/thawed epididymal spermatozoa evaluation

The zona binding capacity of frozen/thawed wisent spermatozoa has been evaluated by using intact heterologous oocytes obtained from frozen/thawed bovine ovaries according to the protocol described before [26] with small modifications.

Control for the ZBA consisted in commercial frozen bull semen of proven fertility.

#### Collection and preparation of the oocytes

Bovine ovaries after obtaining from slaughterhouse were stored at -20°C and re-moved from the freezer 12 hours before planned oocytes collection.

Thawed ovaries were placed on a Petri dishes containing phosphate buffered saline (PBS) and antibiotics (100 IU/mL of penicillin G sodium, 0.1 mg/mL of streptomycin sulfate) with 0.1% w/PVA (PBS/PVA medium) and minced with scalpel blade to release oocytes. Only oocytes with intact zona pellucida were selected by using small bore glass pipette. Selected oocytes were placed in a 4-well dish with PBS/PVA to remove cumulus cells using a 200 μL automatic pipette. Subsequently oocytes were washed twice in modified Tyrode’s-albumin-lactate-pyruvate medium (TALP) [27]. Washed oocytes were placed in 50 ul drops of TALP + penicillamine-hypotaurine-epinephrine (PHE) [28] under paraffin oil (10 oocytes/drop) and put in the incubator (38.5°C in a 5% CO2-air atmosphere (100% relative humidity).

#### Semen preparation

Thawed semen was centrifuged at 500g for 3 minutes and resuspended in TALP+PHE. The subjective motility and concentration was assessed to establish the dilution needed to obtain final concentration of 107 motile sperm/ml. Each drop con-taining oocytes was inseminated with 50 μL of prepared semen. The oocytes were in-cubated with semen for 3 hours at 38.5°C in a 5% CO2-air atmosphere (100% relative humidity).

The zona binding assay was performed in three replicates (n=3). One straw from each epididymis and extender was thawed for each replication. One straw of commercially frozen bull semen was thawed aimed to constitute the control group. The scheme of the experiment is presented in the figure 1.

**Figure 1.**
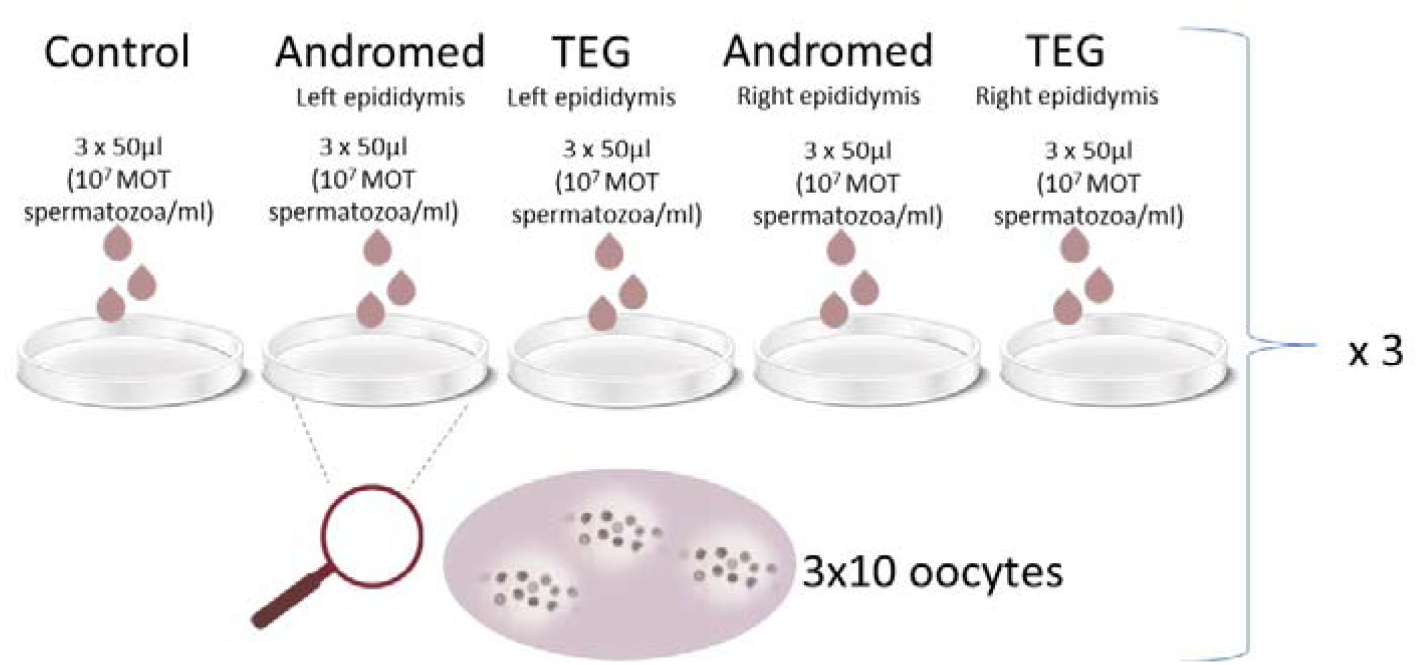
Scheme of the zona pellucida binding assay.

#### Oocyte fixation and staining

After incubation oocyte-sperm complexes were pipetted to remove loosely bound spermatozoa. Subsequently, complexes were fixed with 2.5% glutaraldehyde in PBS for 10 min [29]. After fixation, sperm-oocyte complexes were placed in PBS/PVA in the fridge until the next day.

All sperm-oocyte complexes were evaluated to count the number of adherent spermatozoa by a bis-benzimide (Hoechst 33342) staining. Two small drops of medium with 5 sperm-oocyte complexes each, were placed on the microscope slide and 10 μl of Hoechst Working solution (0.01 mg/mL) was added to each drop and left in the dark for 5 minutes.

After incubation, medium was removed from the slides. 5 μl of antifading solution was added. After 2 minutes, oocytes were covered with cover slip and sealed with silicone.

The number of spermatozoa bound to each oocyte was counted under a fluorescence microscope (x400; Zeiss Axiovert 100).

#### Statistical analysis

The material used in the experiment came from two epididymides of one individual, therefore the results of the spermatozoa analysis were presented as averages and only descriptive statistics were used.

For ZBA results Shapiro-Wilk’s test was used to assess data normality. Nonparametric test (Wilcoxon signed ranks test) were used to evaluate differences between samples. Differences were considered significant at p⍰≤⍰0.05.

An asymptotic test for the equality of coefficients of variation from k populations was used to evaluate the statistical significance of the differences in the results between ZBA replicates [30].

## 3. Results

### 3.1. Characteristics of wisent epididymal spermatozoa obtained post mortem

Subjective motility assessed immediately after collection was assessed as 60% for spermatozoa obtained from both epididymides.

The mean concentration of spermatozoa isolated from the epididymis was 317.5 million/mL.

The mean percentage of viable sperm assessed with the eosin-nigrosin dye was 78.8%.

The mean percentage of sperm with normal morphology was 88.0%.

### 3.2. Basic frozen/thawed semen assessment

Samples frozen/thawed in TEG extender were characterized by higher percentage of motile spermatozoa than those frozen/thawed in Andromed^®^. Which were 25% and 15% respectively. (Fig.2).

**Figure 2.**
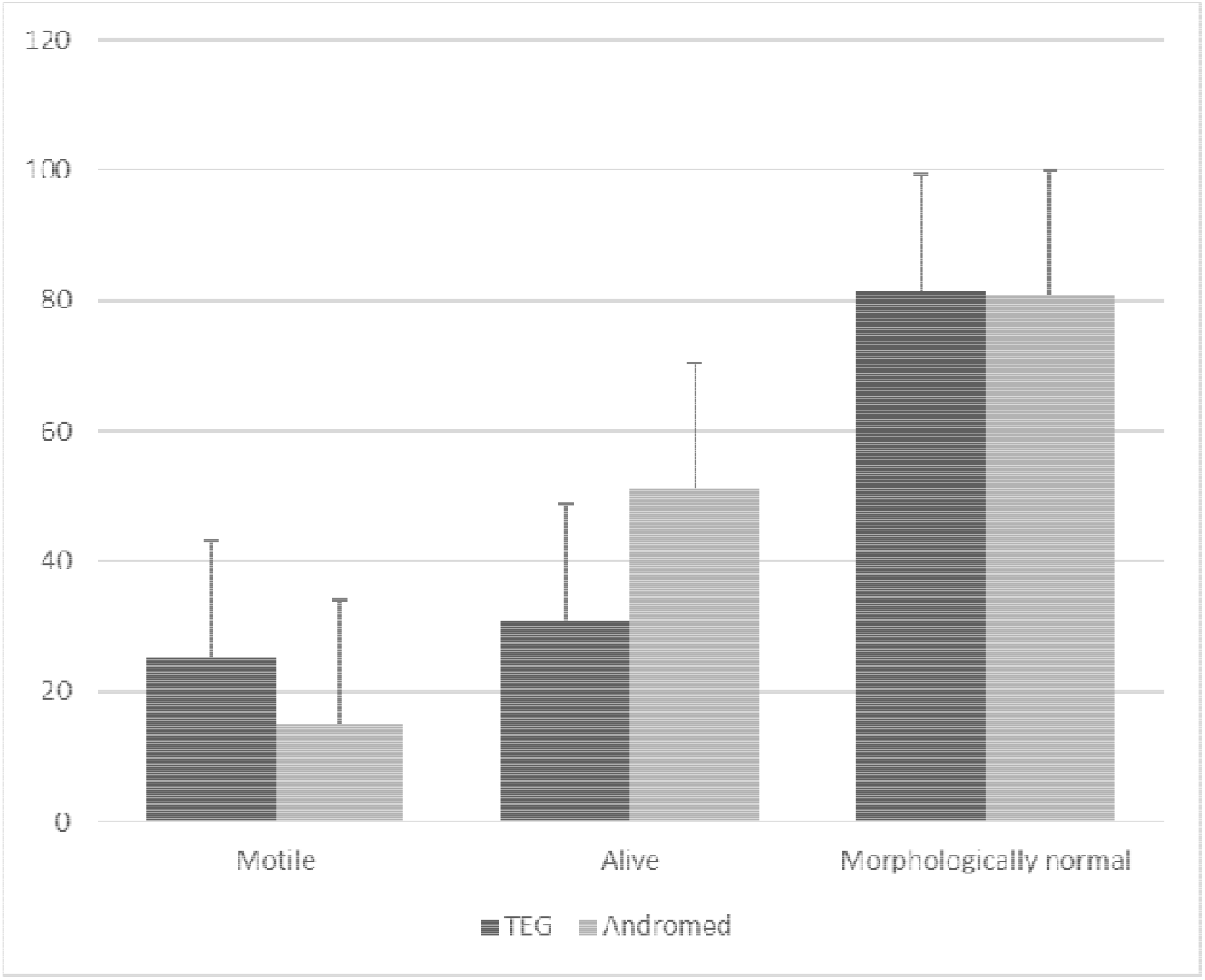
Basic characteristic of frozen/thawed wisent spermatozoa cryopreserved in Tris buffer, hen egg yolk and glycerol based extender (TEG) or Andromed^®^.

The mean percentage of viable spermatozoa, assessed by eosin dye was 30.75 % for TEG and 51.25 % for Andromed^®^ (Fig.2).

The percentage of spermatozoa with normal morphology was comparable for both diluents and amounted to 81.5 % and 80.75 % for TEG and Andromed^®^, respectively (Fig.2).

A comparison of the above-mentioned parameters is shown in Figure 2 (Fig 2).

### 3.3. Characteristic of post-thaw wisent spermatozoa motility assessed by CASA

The motility of the obtained spermatozoa assessed using the CASA system was low for both extenders. The mean percentage of motility (MOT, %) and progressive motility (PMOT, %), were higher in Tris based extender than in Andromed^®^ and were respectively 10% and 3% for TEG and 5.5% and 1% for Andromed^®^. The percentage of the static sperm population was similar in TEG (85.5 %) and Andromed^®^ (83.5 %). In the rapid population, a higher percentage of fast sperm was shown in TEG (5.0 %) than in Andromed^®^ (1.5 %), linearity (LIN) and straightness (STR) were higher in TEG than Andromed^®^ (Figure 3)

**Figure 3.**
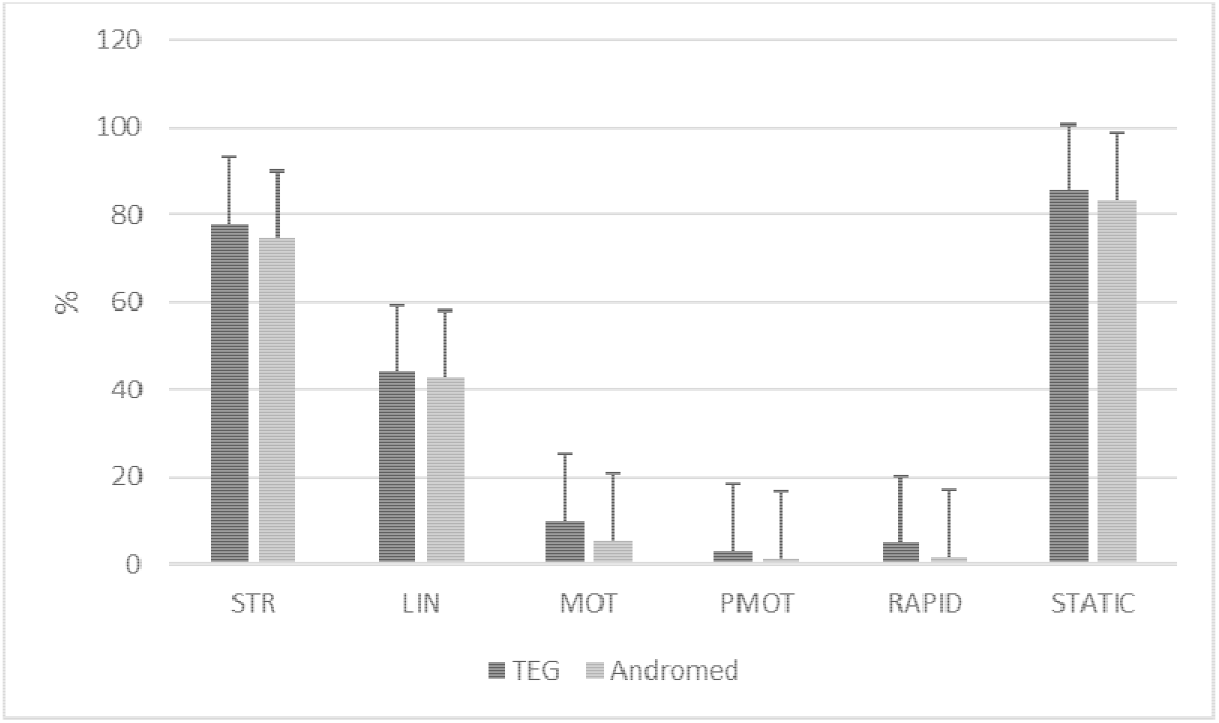
Characteristic of post-thaw wisent spermatozoa cryopreserved with Tris buffer, hen egg yolk and glycerol based extender (TEG) or Andromed^®^ motility assessed by CASA-straightness STR (%); linearity LIN (%); motility MOT (%); progressive motility PMOT (%); percentage of rapid sperm RAPID (%); percentage of static sperm STATIC (%).

The average path velocity (VAP) and straight line velocity (VSL) were higher in TEG. The curvilinear velocity (VCL) was similar in both groups (Figure 4).

**Figure 4.**
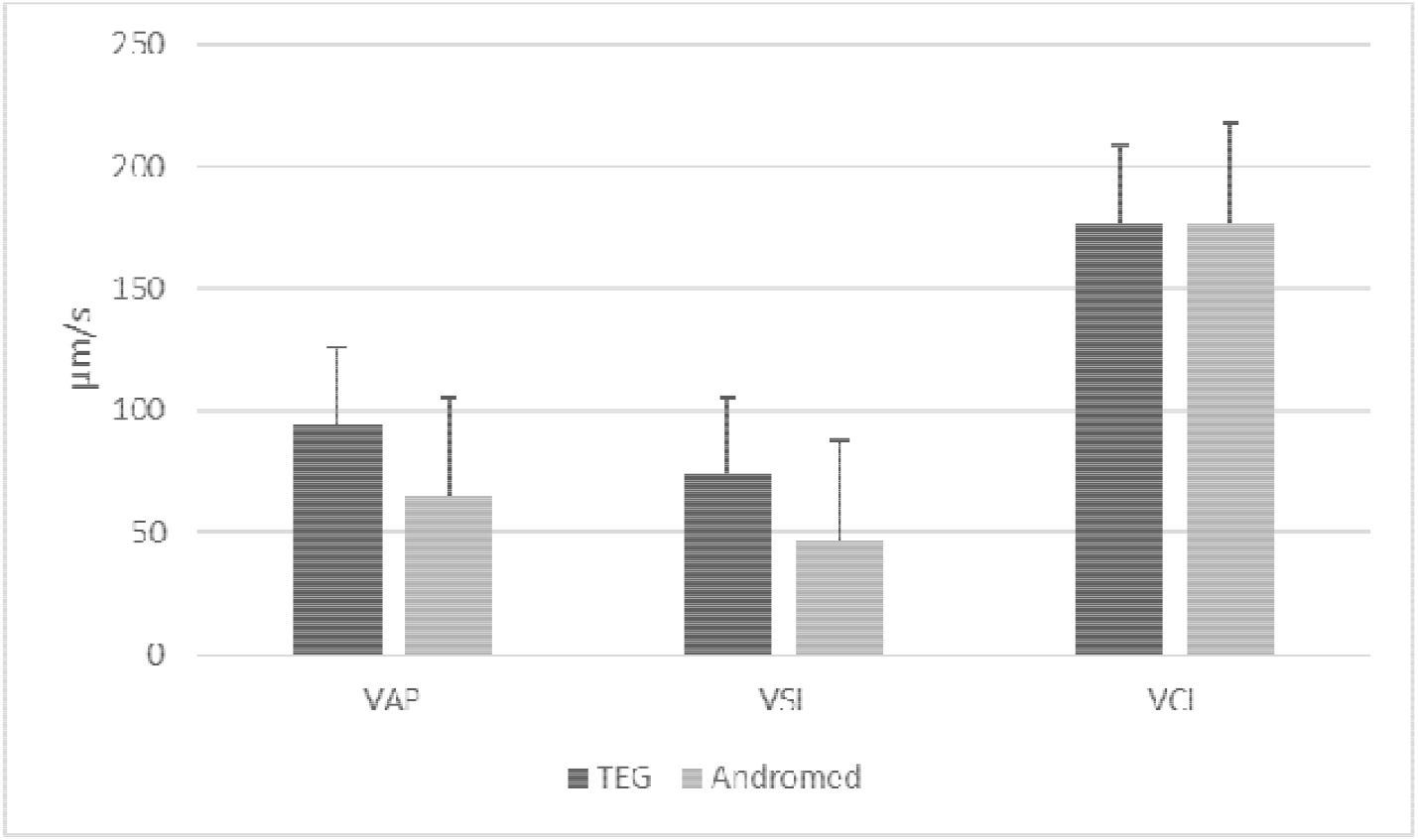
Characteristic of post-thaw wisent spermatozoa cryopreserved with Tris buffer, hen egg yolk and glycerol based extender (TEG) or Andromed^®^ motility assessed by CASA - average path velocity VAP (μm/s); straight line velocity VSL (μm/s); curvilinear velocity VCL (μm/s).

The average Amplitude of lateral head displacement (ALH) was higher in TEG than in Andromed^®^ and amounted to 8.55 μm and 5.1 μm respectively (Figure 5).

**Figure 5.**
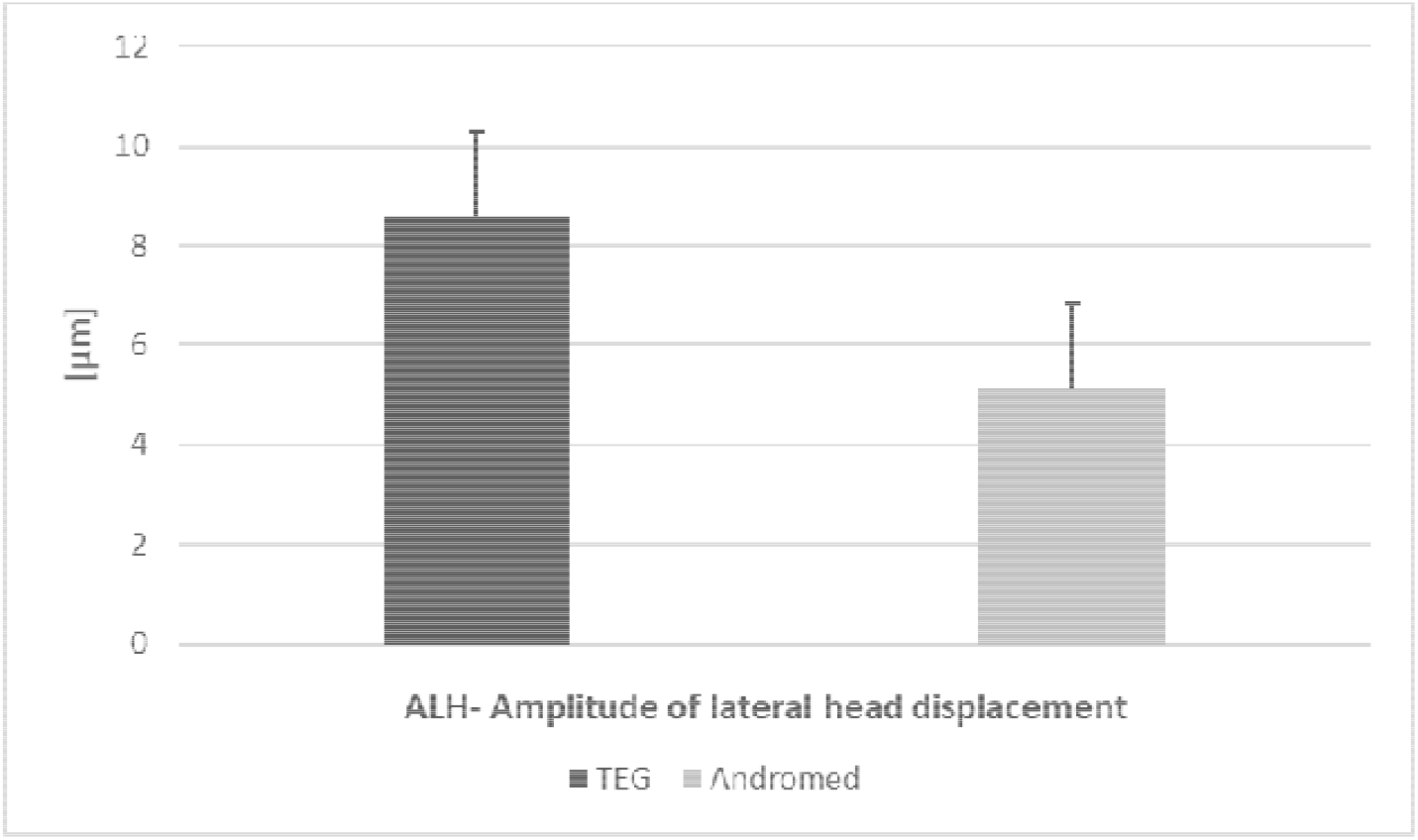
Characteristic of post-thaw wisent spermatozoa cryopreserved with Tris buffer, hen egg yolk and glycerol based extender (TEG) or Andromed^®^ motility assessed by CASA - Amplitude of lateral head displacement ALH (μm)

The beat cross frequency (BCF) was 25.05 Hz for Andromed^®^ and was comparable with 25.00 HZ assessed in TEG (Figure 6).

**Figure 6.**
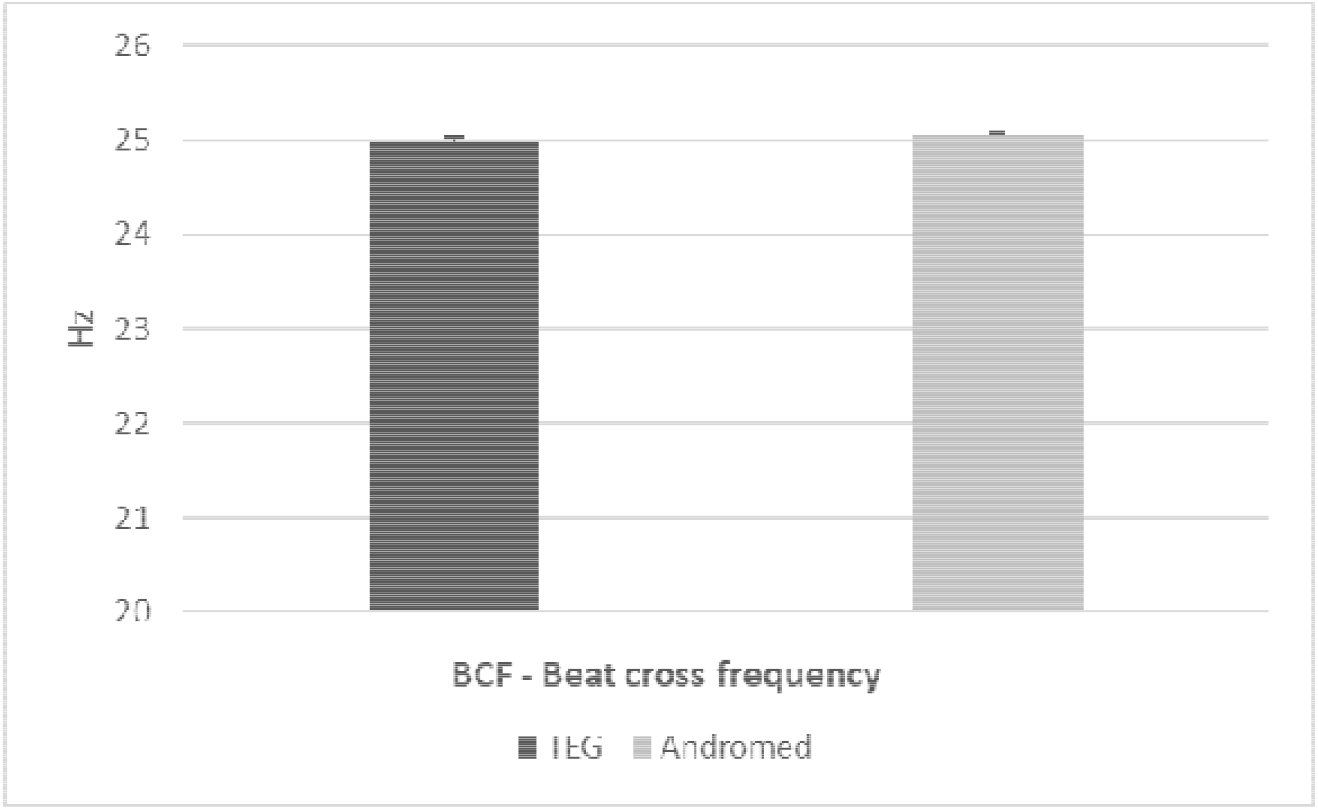
Characteristic of post-thaw wisent spermatozoa cryopreserved with Tris buffer, hen egg yolk and glycerol based extender (TEG) or Andromed^®^ motility assessed by CASA - Beat cross frequency BCF (Hz)

### 3.4 The morpho-functional characteristics of frozen/thawed wisent spermatozoa

The percentage of spermatozoa characterized by an intact sperm membrane and acrosome after thawing was higher in Andromed^®^ than in TEG. Other parameters of morpho-functional characteristics like percentages of viable: nonapoptotic cells, cells with high mitochondrial potential, intact chromatin and without lipid peroxidation were better in samples frozen in TEG. The results of the assessment of functional and structural parameters assessed by flow cytometry are presented in Table 1.

**Table 1.** The functional characteristics of wisent epididymal spermatozoa cryo-preserved in Tris buffer, hen egg yolk and glycerol based extender (TEG) and Andromed^®^. The data are presented as mean. (n=2).

|  | Live cells with intact sperm membrane [%] | Live cells with intact acrosome [%] | Live non apoptotic cells [%] | Live cells with high mitochondrial activity [%] | Live cells with fragmented chromatin [%] | Live cells without lipid peroxidation [%] |
| --- | --- | --- | --- | --- | --- | --- |
| TEG | 14.54 | 29.24 | 46.49 | 34.73 | 0.1 | 17.46 |
| Andromed® | 26.12 | 55.16 | 36.5 | 1.39 | 0.30 | 8.45 |

### 3.5. Zona pellucida binding assay

Regardless of the extender used, all oocytes were with bound spermatozoa. The minimum number of bound spermatozoa was 29 for control, 3 for TEG and 2 for Andromed^®^ groups. The highest number of attached sperm was observed in control group for all replicates. The mean number of bound spermatozoa was higher in extender based on Tris and chicken egg yolk than in Andromed^®^, but in both extenders it was markedly lower than in control (Table 2). Representative images are reported in Figure 7 and 8.

**Table 2.** Post-thaw ability of wisent epididymal spermatozoa cryopreserved in Tris buffer, hen egg yolk and glycerol based extender (TEG) and Andromed^®^ to bind heterologous zonae pellucidae. Data are presented as mean ± SE of 6 replicates (3 for samples from left epididymis and 3 from right), 30 oocytes per replicate in each group. a,b, care significantly different (p⍰<⍰0.05)

|  | The mean number<br>of spermatozoa<br>bound to oocyte | Oocytes with bound<br>spermatozoa (%) |
| --- | --- | --- |
| Control | 166.11 ± 50.60 | 100.00 |
| TEG | 66.26 ± 7.93 | 100.00 |
| Andromed® | 27.37 ± 2.37 | 100.00 |

**Figure 7.**
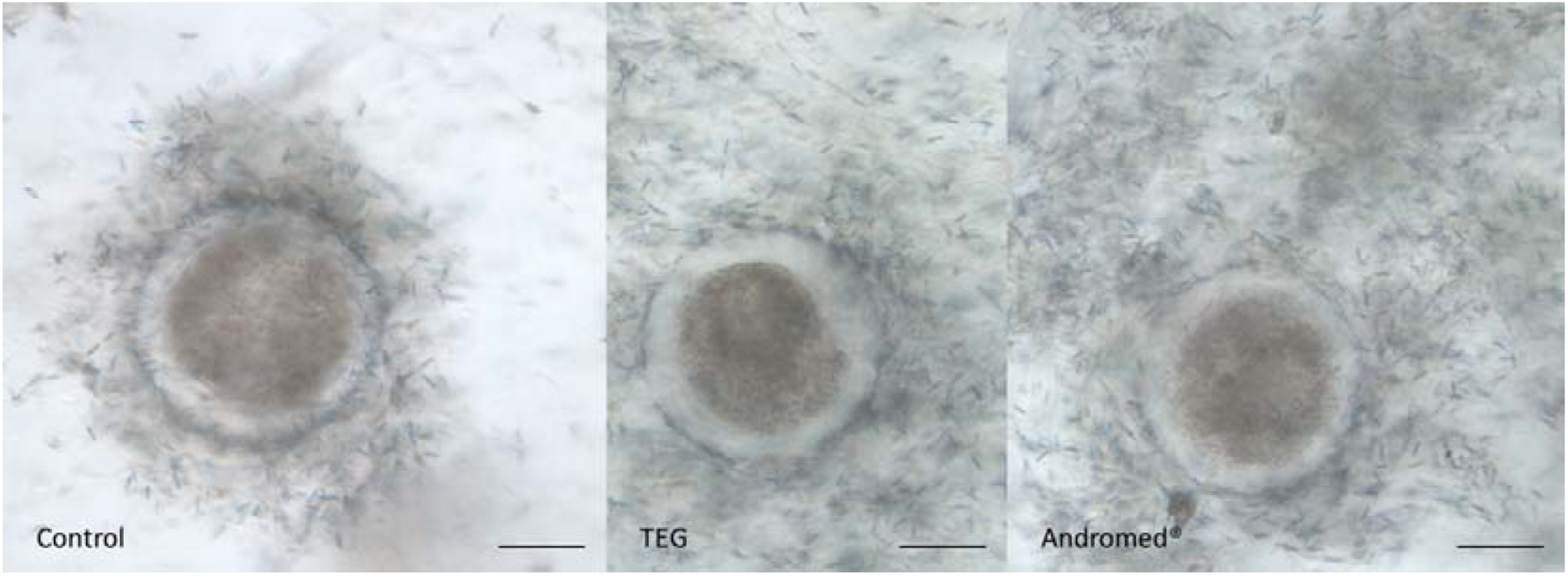
Representative pictures of Zona Binding Assay of wisent spermatozoa cryopreserved with two different extenders (TEG (Tris buffer, egg yolk and glycerol based extender), or Andromed ^®^) and control bull cryopreserved semen. Objective 40X, scale bar = 50μm.

**Figure 8.**
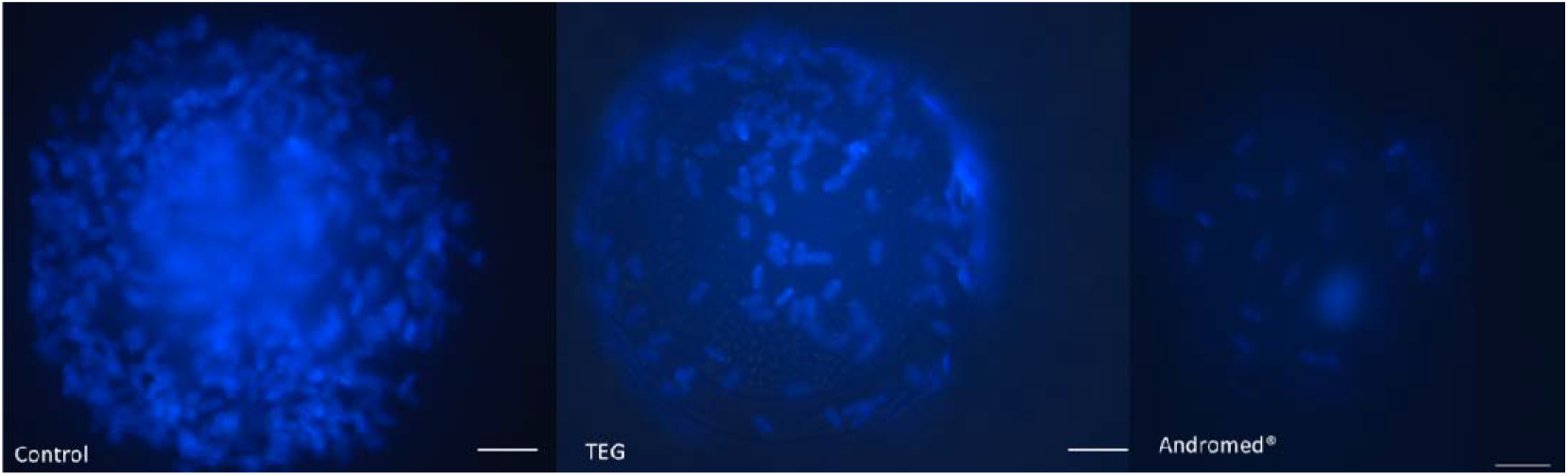
Representative pictures of Zona Binding Assay of wisent spermatozoa cryopreserved with two different extenders (TEG (Tris buffer, egg yolk and glycerol based extender), or Andromed ^®^) and control bull cryopreserved semen. Hoechst 33342 staining. Objective 40X, scale bar = 20 μm.

The value of the test statistic in the asymptotic test of homogeneity of coefficients of variation was 58.74 (p=0.85). The differences in the variability of the results for 6 repetitions are not statistically significant.

## 4. Discussion

Epididymal spermatozoa collected post mortem are considered as a valuable source of gametes for protected animals genome banks [4]. For some species, such as European bison, they are the only source of male gametes that can be cryopreserved and used later in ART [2,8]. The possibility of effective cryopreservation of wisent epididymal spermatozoa has been presented in a few publications [8,9]. However, to be able to fully utilize their potential, deep knowledge of them is needed [1,4]. Spermatozoa collected post mortem from a protected species are extremely valuable, but also problematic research material, due to limitation in their availability. Therefore, there is a need do make the choice for which research to devote them. It is desirable, to keep the sufficient amount of spermatozoa for ART and at the same time obtain as much information as possible about them. For this reason, research on methods for their full post thawed characterization are being still developed. Cases of cryopreservation of wisent epididymal spermatozoa and the assessment of its effects described so far in the literature, were grounded on basic methods (viability, subjective motility and morphology), computer-assisted sperm analysis and flow cytometry [8,9].

It was observed that defective acrosome reaction and/or zona pellucida interaction were frequently noticed in the semen of infertile man, regardless of the presence of normal or abnormal results of basic sperm assessment [11]. Semen assessment with the use of the CASA system makes it possible to assess the quality of the spermatozoa obtained on the basis of the detailed characteristics of their movement. As it was described in cattle, semen samples with large population of progressive and rapid spermatozoa are prone to have better sperm longevity after thawing [4]. It was described that velocity parameters such as average path velocity (VAP) and straight line velocity (VSL) are positively correlated with post-thaw motility, thus could constitute potential bull fertility markers [4]. However, there are too many factors, such as variation in semen parameters between individuals and ejaculates from the same individual, and the conditions of transport, storage and dilution of semen, that can negatively affect the results of evaluation by the CASA system.

Flow cytometry is a method that allows the evaluation of morpho-functional features of the spermatozoa. The standard flow cytometry protocol for semen evaluation used by the authors includes the evaluation of such features determining proper functioning of spermatozoa as: cell membrane and acrosome integrity, lipid peroxidation, apoptosis like changes, mitochondrial activity, membrane lipid disorders and chromatin status. The results of each of these parameters allow some conclusions to be drawn regarding the fertility of the collected and stored samples. The findings of each of these methods provide information about the quality of the semen obtained, but none of them alone can determine its fertilizing potential.

The fertilization potential of frozen wisent epididymal spermatozoa was evaluated in vivo tests by insemination of Holstein Friesian heifers. As a result of the cited study, domestic cattle and wisent hybrids were born [9]. With the use of gametes frozen in the Tris, egg yolk and glycerol based extender (TEG), an European bison early blastocyst was obtained in vitro [21]. The mentioned studies prove the fertilization capacity of European bison spermatozoa collected and cryopreserved in this way. However, there are no procedures described so far that would allow for the classification and evaluation of the fertilization potential of the stored material. For the protection of species diversity, the significant value of each collected genetic material is undeniable [2]. However, being able to approximately assess its fertilization potential, we can decide on its intended use - whether sperm frozen in a given collection can be used in the future for artificial insemination or whether their potential is sufficient to carry out classic in vitro ART (e.g., IVF).

The zona pellucida binding assay makes it possible to assess the sperm’s ability to attach to the zona pellucida [14]. Binding of the sperm with the oocyte is the basic and one of a series of stages leading to fertilization. It takes place through the connection of a receptors on the spermatozoa membrane and glycoproteins of zona pellucida [22]. Therefore, this method allows to evaluate sperm for disturbances in the structure of the membrane that are not detectable by routine semen analysis procedures [14]. The correlation between ZBA results and in vivo fertility was observed in cattle [15]. The ability to bind to the oocyte is a characteristic features of living cells [13-15]. The obtained results of viability and sperm membrane integrity seem to confirm this also in European bison.

The ZBA protocol used, based on research conducted on a related species, turned out to be suitable for the assessment of wisent epididymal spermatozoa. Both in the control group and in the groups of studied extenders, spermatozoa attached to all oocytes. The quality of cryopreserved spermatozoa obtained after thawing was low regardless of the extender used. It is reflected in the results of their assessment using basic methods, CASA and flow cytometry, which support the results of ZBA.

Taking into account basic parameter of semen assessment-motility after thawing, samples from the control group were characterized by a higher percentage of subjective motility than in the both study groups. The results of detailed movement characterization using the CASA system showed better results of sperm frozen in TEG than in Andromed^®^. In addition, the mitochondrial potential assessed by the flow cytometer supports the CASA results. Therefore, the motility seems to match the results obtained in ZBA, although it should be remembered that the ZBA protocol was performed based on the same concentration of motile gametes in all the groups. The percentage of spermatozoa characterized by an intact sperm membrane and acrosome after thawing was higher in Andromed^®^ than in TEG. Other parameters of structural and functional features and CASA were better in samples frozen in TEG. Ultimately, the results of the ZBA test seem to speak in favor of the TEG extender. This suggests that the analyzes performed so far are not able to detect particular membrane defects that affected the results of ZBA. However, following the sperm binding to the zona pellucida, acrosomal reaction requires a properly constructed acrosome [23]. For this reason, it could turn out that Andromed^®^ could show better results in the zona pellucida penetration test or IVF. However, this requires repeating the comparison of diluents on a larger number of trials. Continuing this extenders comparison in the future, the variability between individuals should also be taken into account and the study group should be enlarged.

In cattle, a large variability in sperm-binding capacity has been observed within replicates and between ZBA replicates, which could also be due to oocyte variability [13]. In the presented experiment, although oocytes from multiple ovaries were used in order to reduce variability, we also noticed this variation between repetitions. However, regardless of the repetition, each time the sperm frozen in TEG showed a better ZBA result compared to those frozen in Andromed^®^. However, the use of two extenders was aimed to authenticate the assessment of the usefulness of the applied ZBA protocol for testing wisent epididymal spermatozoa. Therefore, the results described in the presented article cannot unambiguously assess the advantage of one of the extenders.

## 5. Conclusions

It is not possible to evaluate the actual fertilization capacity of spermatozoa on the basis of the tests carried out so far in European bison. However, the zona pellucida binding assay is a step further in evaluating the reproductive potential of the collected material. The ZBA protocol using frozen bovine oocytes, developed for bull semen, is relatively simple to perform. Carried out experiment showed that it is suitable for European bison and therefore, should be included in the routinely used protocol for the evaluation of cryopreserved epididymal sperm from wisent.

## Author Contributions

M.E.—conceptualization, performing the study, writing the manuscript; M.C. —performing and supervising the study, editing the manuscript; S.P.— correcting and editing the manuscript; G.C.L.— reviewing and correcting manuscript; W.O.—reviewing and correcting, funding acqui-sition; W.N.— conceptualization, reviewing and correcting, funding acquisition

## Funding

This research was funded by the Forest Fund (Poland), grant number OR.271.3.10.2017 and supported by the Polish National Agency for Academic Exchange under Grant No. PPI/APM/2019/1/00044/U/00001.

Institutional Review Board Statement: No animal was killed to obtain material for these studies. The Approval of the local ethics committee was not required.

## Data Availability Statement

The data that support the findings of this study are available from the corresponding authors [ME &WN], upon reasonable request.

